# OrganoID: a versatile deep learning platform for tracking and analysis of single-organoid dynamics

**DOI:** 10.1101/2022.01.13.476248

**Authors:** Jonathan Matthews, Brooke Schuster, Sara Saheb Kashaf, Ping Liu, Rakefet Ben-Yishay, Dana Ishay-Ronen, Le Shen, Christopher Weber, Margaret Bielski, Sonia S. Kupfer, Mustafa Bilgic, Andrey Rzhetsky, Savaş Tay

## Abstract

Organoids have immense potential as *ex vivo* disease models for drug discovery and personalized drug screening. Dynamic changes in individual organoid morphology, number, and size can indicate important drug responses, however these metrics are difficult and labor-intensive to obtain for high-throughput image datasets. Here, we present OrganoID, a robust image analysis platform that automatically recognizes, labels, and tracks single organoids, pixel-by-pixel, in brightfield and phase-contrast microscopy experiments. The platform was trained on images of pancreatic cancer organoids and validated on separate images of pancreatic, lung, colon, and adenoid cystic carcinoma organoids, which showed excellent agreement with manual measurements of organoid count (96%) and size (95%) without any parameter adjustments. Single-organoid tracking accuracy remained above 89% over a four-day time-lapse microscopy study. Automated single-organoid morphology analysis of a chemotherapy dose-response experiment identified decreased organoid circularity as an important morphological feature reflecting drug response. OrganoID enables straightforward, detailed, and accurate image analysis to accelerate the use of organoids in high-throughput, data-intensive biomedical applications.

## INTRODUCTION

Three-dimensional (3D) cell culture systems serve as more physiologically relevant *in vitro* models than traditional human cell monolayers for both basic and clinical applications^1^. Organoids are multicellular 3D structures derived from primary or stem cells that are embedded into a biological matrix to create an extracellular environment that provides structural support and key growth factors. Organoids are particularly useful models as they closely recapitulate cellular heterogeneity, structural morphology, and organ-specific functions of a variety of tissues^2–5^. Live-cell imaging can reveal important features of organoid dynamics, such as growth, apoptosis/necrosis, and movement, which can reflect physiological and pathological behaviors and drug responses. While organoids have been successfully used to investigate important phenomena that might be obscured in simpler models, their use in data-intensive applications, such as high-throughput screening, has been more difficult.

A major challenge for organoid experiments is drug-response measurement and analysis, which must be performed for a large number of microscopy images. Image analysis is particularly difficult for organoid experiments due to their movement across focal planes and variability in organoid size and shape between different tissue types, within the same tissue type, and within the same single culture sample^6,7^. Several recently developed organoid platforms, while powerful, rely on per-experiment or per-image tuning of brightfield image analysis parameters^8–10^, or require manual labeling of each image^11–13^, which limits experiment reproducibility and scale. Organoids can be fluorescently labeled to aid in image segmentation and tracking, such as through genetic modification for fluorescent protein expression^14–16^, cell fixation and staining^8,15^, or the use of membrane-permeable dyes. However, these approaches may alter intrinsic cellular dynamics from original samples^17,18^, limit measurements to endpoint assays, or cause cumulative toxicity through longer growth times and limited diffusion through the hydrogel matrix^19^. There is a critical need for an automated image analysis tool that can robustly and reproducibly measure live-cell organoid responses in high-throughput experiments^9-13^ without the use of potentially toxic or confounding fluorescence techniques.

A number of software tools have been developed to automate the process of brightfield/phase-contrast organoid image analysis. These platforms use conventional image processing methods, such as adaptive thresholding and mathematical morphology^20^, or convolutional neural networks^21–23^ to identify organoids in sequences of microscopy images. Despite their advantages, many existing platforms require cellular nuclei to be transgenically labeled^22^, which increases experiment time and complexity and may modify cellular dynamics. Other existing platforms require manual tuning of parameters for each image^20^, focus on single-timepoint analysis^23^, only provide population-averaged (bulk) measurements without single-organoid resolution, or are limited to bounding-box detection^21^, which fails to capture potentially useful morphological information at the individual organoid level. Changes in organoid shape, such as spiking or blebbing, can reveal important responses to external stimuli and might be missed with bounding-box measurements^24^. Many of these existing platforms were also developed for analysis of organoids derived from a single type of tissue and for images obtained with one specific geometric configuration, precluding their use across different experimental settings.

To address these challenges, we have developed a software platform, OrganoID, that can identify and track individual organoids in a population derived from a wide range of tissue types, pixel by pixel, in both brightfield and phase-contrast microscopy images and in time-lapse experiments. OrganoID consists of (i) a convolutional neural network, which detects organoids in microscopy images, (ii) an identification module, which resolves contours to label individual organoids, and (iii) a tracking module, which follows identified organoids in time-lapse imaging experiments. Most importantly, OrganoID was able to accurately segment and track a wide range of organoid types, including those derived from pancreatic ductal adenocarcinoma, adenoid cystic carcinoma, and lung and colon epithelia, without cell labeling or parameter tuning. The OrganoID software overcomes a major hurdle to organoid image analysis and supports wider integration of the organoid model into high-throughput applications.

## RESULTS

### A robust convolutional neural network for pixel-by-pixel organoid detection

We developed a deep learning-based image analysis pipeline, OrganoID, that recognizes and tracks individual organoids, pixel by pixel, for bulk and single-organoid analysis in brightfield and phase-contrast microscopy images **(Figure 1a)**. The platform employs a convolutional neural network to transform microscopy images into organoid probability images, where brightness values represent the network belief that an organoid is present at a given pixel. The network structure was derived from the widely successful *u-net* approach to image segmentation^25^ **(Figure 1b)**. The *u-net* approach first passes each image through a contracting series of multidimensional convolutions and maximum filters to extract a set of deep feature maps that describe the image at various levels of detail and contexts, such as edges, shapes, and textures. The feature maps are then passed through an expanding series of transposed convolutions, which learn to localize the features and assemble a final output. The OrganoID neural network follows a *u-net* structure and was optimized to require far fewer feature channels than the original implementation. Model simplification can limit overfitting and minimizes the amount of memory and computational power required to use the network in the final distribution. Additionally, all hidden convolutional layers were set to compute outputs with the exponential linear unit (ELU) activation function^26^, which has demonstrated higher accuracy than the rectified linear unit (ReLU) and avoids vanishing gradient problems during network training. Neurons in the final convolutional layer were set with a sigmoid activation function to produce a normalized output that corresponds to the probability that an organoid is present at each pixel in the original image.

**Figure 1.**
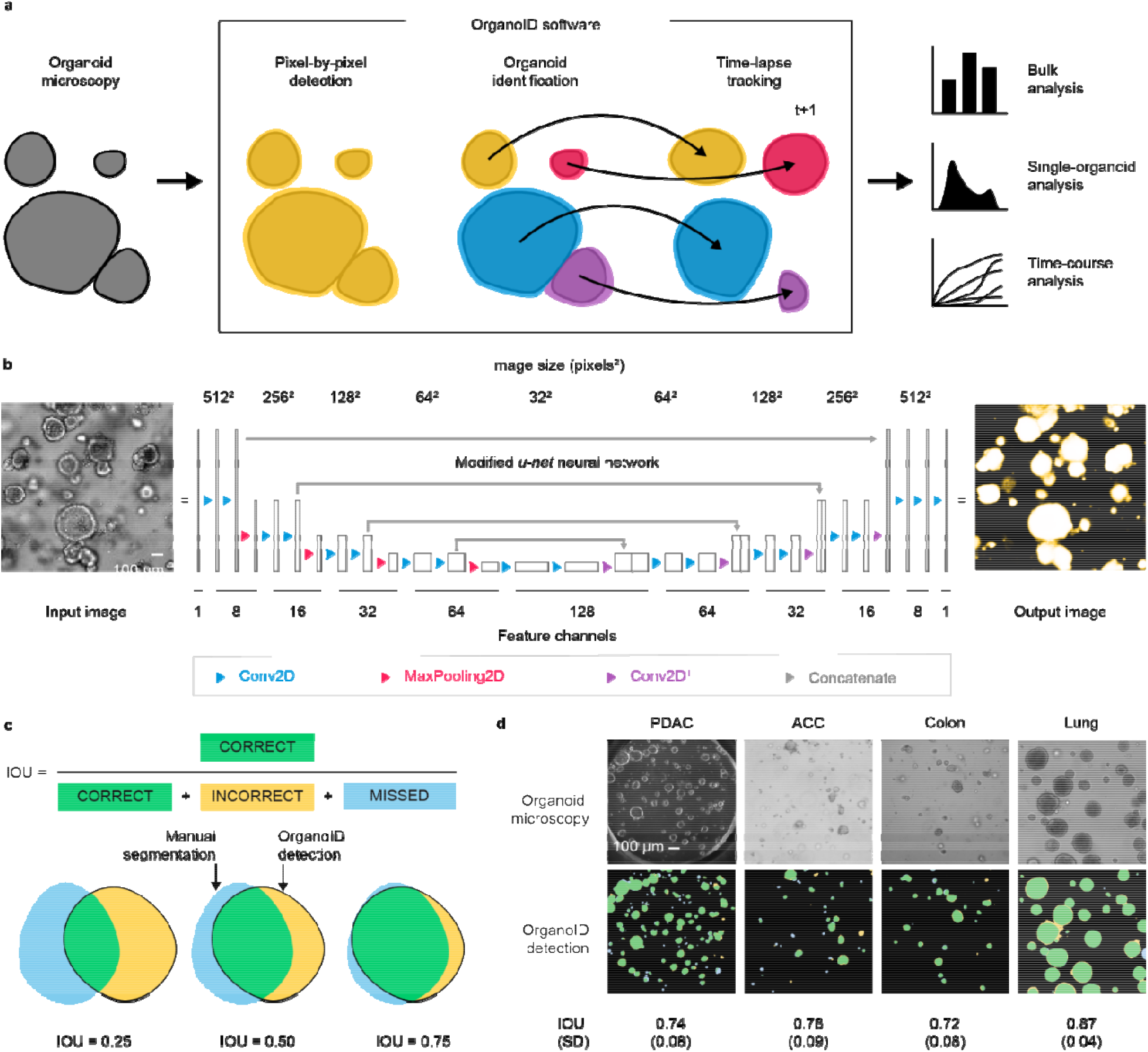
**(a)** The OrganoID software automates robust analysis of organoid microscopy images. Contours are detected pixel by pixel and then separated into distinct organoids for bulk or single-organoid analysis. Identified organoids can also be tracked across time-lapse image sequences to follow responses over time. **(b)** Microscopy images are processed by a convolutional neural network to produce images that represent the probability that an organoid is present at each pixel. The network follows the *u-net* architecture, which applies a series of two-dimension convolutions, maximum filters, and image concatenations to extract and localize image features. Feature channel depths were minimized to limit overfitting and computational power required to use the tool. Scale bar 100 µm. **(c)** The intersection-over-union (IOU) metric, defined as the ratio of the number true positive pixels to the union of all positive pixels, was used to measure the quality of the neural network detections. To compute the IOU, pixels above 0.5 in the network prediction image were marked as positives. Examples of IOU values for several degrees of overlap are shown. **(d)** A representative set of images of different organoid types from the test dataset are shown with the corresponding OrganoID detections. Detections are overlayed on top of ground truth measurements and colored according to the schema in (c). The mean and standard deviation of IOU for images of each organoid type in the test dataset are also shown. Scale bar 100 µm.

An original dataset of 66 brightfield and phase-contrast microscopy images of organoids were manually segmented to produce black-and-white ground truth images for network training and validation **(Supp. Figure 1a)**. Each image featured 5 to 50 organoids derived from human pancreatic ductal adenocarcinoma (PDAC) samples from two different patients. Organoids were either grown on a standard tissue culture plate or on a microfluidic organoid platform^27^. To teach the network that segmentations should be independent of imaging orientation, field-of-view, lens distortion, and other potential sources of overfitting, the training dataset was augmented with random rotation, zoom, elastic distortion, and shear transformations to produce a total of 2,000 images **(Supp. Figure 1b)**.

Network training halted after 37 epochs, once segmentation error (binary cross-entropy) on the validation dataset converged to a minimal value **(Supp. Figure 1c, Video 1)**. After hyperparameter tuning, the final model performance was assessed on a novel PDAC organoid testing dataset, previously unseen by the network. Performance was quantified with the intersection-over-union (IOU) metric, which is defined as the overlap between the predicted and actual organoid pixels divided by their union **(Figure 1c)**. An IOU greater than 0.5 is generally considered to reflect a good prediction and we chose this value to be the minimal benchmark for satisfactory model performance. All PDAC images passed the benchmark, with a mean IOU of 0.74 (SD = 0.081) **(Supp. Figure 2)**.

Because the network was trained and assessed solely with images of PDAC organoids, we were also curious to evaluate its capacity to generalize to organoids derived from non-PDAC tissues. 18 microscopy images of organoids derived from lung epithelia, colon epithelia, and salivary adenoid cystic carcinoma (ACC) were manually segmented. The OrganoID network passed the benchmark for all non-PDAC images, with a mean IOU of 0.79 (SD=0.096). These results demonstrate the capability of the OrganoID neural network to reliably detect organoids from various tissues of origin, from both malignant and non-malignant tissues, in a pixel-by-pixel manner **(Figure 1d)**.

The network was also evaluated for appropriate exclusion of non-organoid technical artifacts. Air bubbles in culture media or gel matrix were rarely detected by OrganoID with a false positive rate of 4.2% **(Supp. Figure 3a)**. We observed that OrganoID segmentations also avoided cellular debris or dust embedded into the gel, ignored chamber borders, and performed robustly across microscope resolutions, organoid concentrations, and organoid shapes **(Supp. Figure 3b-g)**. OrganoID proved to be computationally efficient; each image was segmented in ∼300 milliseconds on a laptop CPU (Intel i7-9750H, 2.6GHz) with less than 200 megabytes of RAM usage.

### Identification of individual organoids with diverse morphology and size

The convolutional neural network detects organoids in an image on a pixel-by-pixel basis, which can be used to measure bulk responses. For single-organoid analysis, pixels must be grouped together to identify individual organoids. This task is straightforward for isolated organoids, where all high-belief pixels in a cluster correspond to one organoid, but is more difficult for organoids that are in physical contact. To address this challenge, we developed an organoid separation pipeline **(Figure 2a)** that uses the raw network prediction map to group pixels into single-organoid clusters. Conventionally, neural network image segmentation methods set an absolute threshold on predicted pixels to produce a binary detection mask. This approach is effective, but discards useful information about the strength of predictions. The segmentations that were used to train the neural network were produced with a thin boundary between organoids in contact. As a result, the network predictions were marginally less confident about pixels near organoid boundaries. We took advantage of this phenomenon to identify and separate organoid contours with a modified Canny edge detector^28^ and a watershed transformation. Our detector sequentially applies (i) a pair of Sobel operators, which compute the image intensity gradient; (ii) a Gaussian filter, which smooths noisy regions; and (iii) a hysteresis-based threshold, which identifies locally strong edges. Edges are removed from the thresholded prediction image to mark the centers of each organoid. These centers are then used as initializer basins for a watershed transformation applied to the raw prediction image. The image is further refined to remove organoids that may be partially out of the field-of-view or are below a particular size threshold. The pipeline outputs a labeled image, where the pixels that represent an individual organoid are all set to a unique organoid ID number, which can be used for single-organoid analysis **(Figure 2a)**.

**Figure 2.**
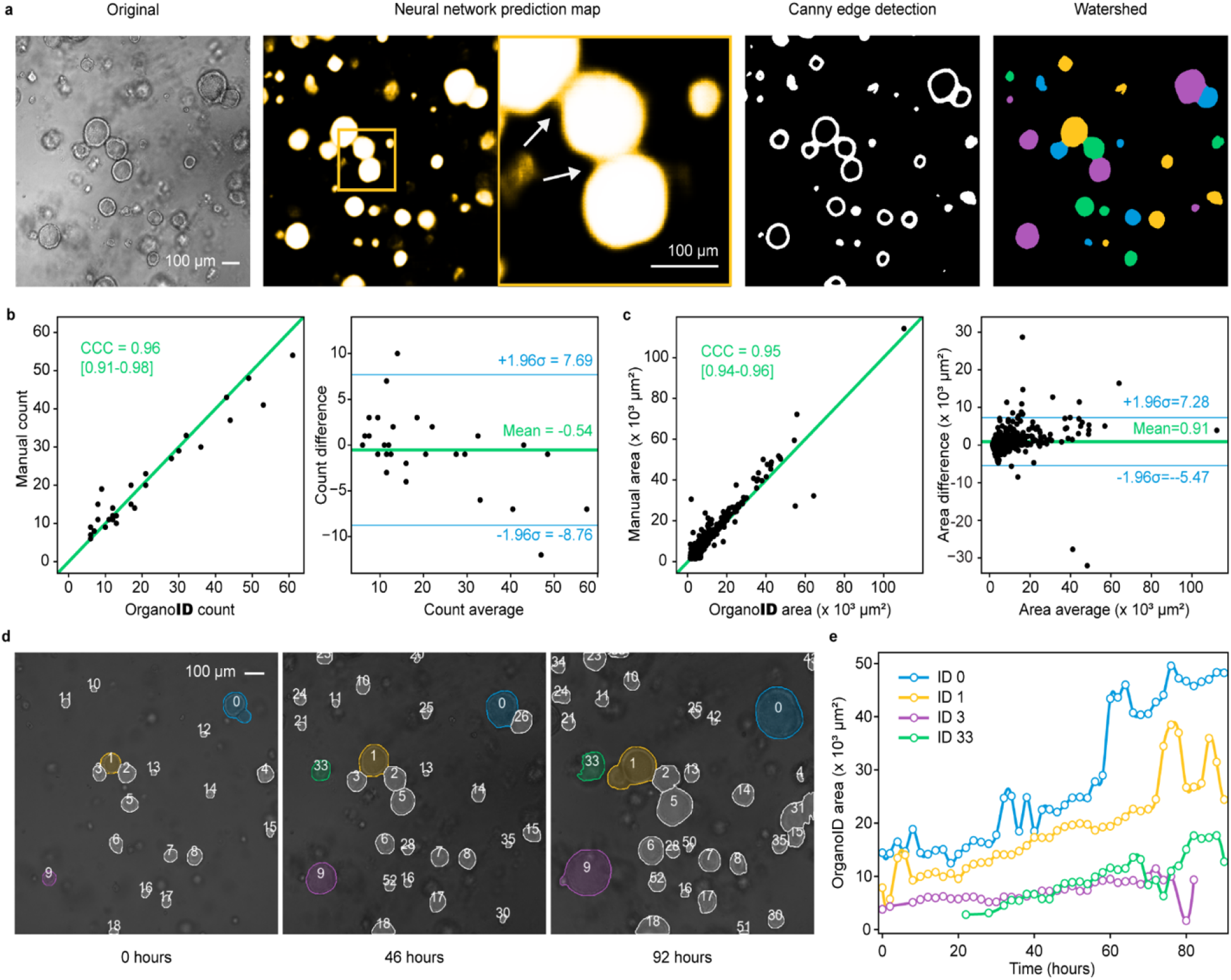
**(a)** OrganoID can identify individual organoid contours, including those in physical contact. An example image (left) is shown to demonstrate the steps of single-organoid identification. The neural network predictions (second-from-left) are observably less confident for pixels at organoid boundaries (enlarged view, indicated with white arrows), which enables edge detection with a Canny filter (second-from-right). Edges are used to identify organoid centers, which serve as basin initializers for a watershed transformation on the prediction image to produce a final single-organoid labeled image (right). **(b)** The identification pipeline was used to count the number of organoids in images from the test dataset. These counts were compared to the number of organoids in the corresponding manually segmented images. The concordance correlation coefficient (CCC) was computed to quantify measurement agreement (left). Bland-Altman analysis (right) demonstrates low measurement bias and limits of agreement. Black dots are test images. Green line in the left plot is *y=x*. **(c)** The area of each organoid in all test images was also measured manually and with OrganoID. Measurements were compared with CCC computation (left) and Bland-Altman analysis (right). Black dots are identified organoids. **(d)** Identified organoids in time-lapse microscopy images are matched across frames to generate single-organoid tracks and follow responses over time. Shown are images of three timepoints from an organoid culture experiment. **(e)** Automatically measured growth curves for a selected set of organoids from the experiment in (d).

For quantitative validation, OrganoID was used to count and measure the area of organoids in images from the PDAC and non-PDAC testing datasets (a total of 28 images). These data were then compared to the number and area of organoids in the corresponding manually segmented images. Organoid counts agreed with a concordance correlation coefficient (CCC) of 0.96 [95% CI 0.91-0.98]. OrganoID, on average, detected 0.5 fewer organoids per image than manual segmentation. The limits of agreement between OrganoID and manual counts were between -8.5 and 7.5 organoids (**Figure 2b**). Organoid area comparison demonstrated a CCC of 0.95 [95% CI 0.94-0.96]. OrganoID area measurements were biased to be 0.91×10^3^ μm^2^ larger per organoid with limits of agreement between -5.47×10^3^ μm^2^ and 7.28×10^3^ μm^2^ (**Figure 2c**). Overall, the measurements produced by OrganoID were in considerable agreement with those obtained by hand, which supports the use of OrganoID for automated single-organoid analysis.

### Automated tracking of individual organoids in time-lapse microscopy experiments

OrganoID was also developed with a tracking module for longitudinal single-organoid analysis of time-course imaging experiments, where changes in various properties, such as size and shape, of individual organoids can be measured and followed over time. The central challenge for the tracking module is to match a detected organoid in a given image to the same organoid in a later image. The Hungarian assignment algorithm^29^ was used to minimize a cost matrix based on the number of shared pixels between detected organoids in two images. This approach produces organoid “ tracks” for unique organoids in time-lapse images. For validation, microscopy images taken every 4 hours from a 92-hour organoid culture experiment were passed through the entire OrganoID pipeline to produce growth curves that followed single-organoid changes over time **(Figure 2d, e)**. The tracking step was also performed by hand to evaluate automated performance, which maintained over 89% accuracy throughout the duration of the experiment **(Supp. Figure 4, Video 2)**.

### OrganoID measures bulk and single-organoid death responses over time

PDAC organoids were treated with serial dilutions of gemcitabine (3nM-1000nM), an FDA-approved chemotherapeutic agent commonly used to treat pancreatic cancer, and imaged every 4 hours of the course of 72 hours. Propidium iodide (PI), a fluorescent reporter of cellular necrosis, was also added to the culture media to monitor death responses. Brightfield images were then processed with the OrganoID platform to identify organoids and analyze bulk and single-organoid responses **(Figure 3a)**. At gemcitabine concentrations above 3 nM, the total organoid area increased for the first several hours, which reflected initial organoid growth, but then decreased to a value and at a rate inversely proportional to gemcitabine concentration **(Figure 3b)**. Identified organoid counts for gemcitabine concentrations above 10 nM also sharply decreased over time. The total fluorescence intensity of the PI signal increased over time to a value and at a rate proportional to gemcitabine concentration, however the response to 100 nM gemcitabine appeared to induce a stronger death signal than the response to 1000 nM gemcitabine **(Figure 3c)**. Measurements of death stains such as PI are typically normalized to a viability measurement that accounts for differences in the number and size of organoids between replicates that can compound over the duration of an experiment. We used OrganoID to normalize live-cell fluorescence measurements: the total fluorescence intensity of identified organoid regions was divided by the total organoid area at each time point. Normalization with OrganoID increased the separation of responses between each treatment group over time, decreased standard error across replicates, and corrected the response discrepancy between the 100 nM and 1000 nM conditions **(Figure 3c)**. We also compared these normalized measurements to normalization with an MTS proliferation assay, which can only be used for endpoint analysis. Normalization with OrganoID identified the same significantly different endpoint responses as MTS assay normalization **(Supp. Figure 5, Supp. Table 1)**. The fluorescence of individual organoids did not significantly differ across concentrations of gemcitabine **(Supp. Figure 6)**, however fluorescence normalized to single-organoid area was significantly different (p<0.001).

**Figure 3.**
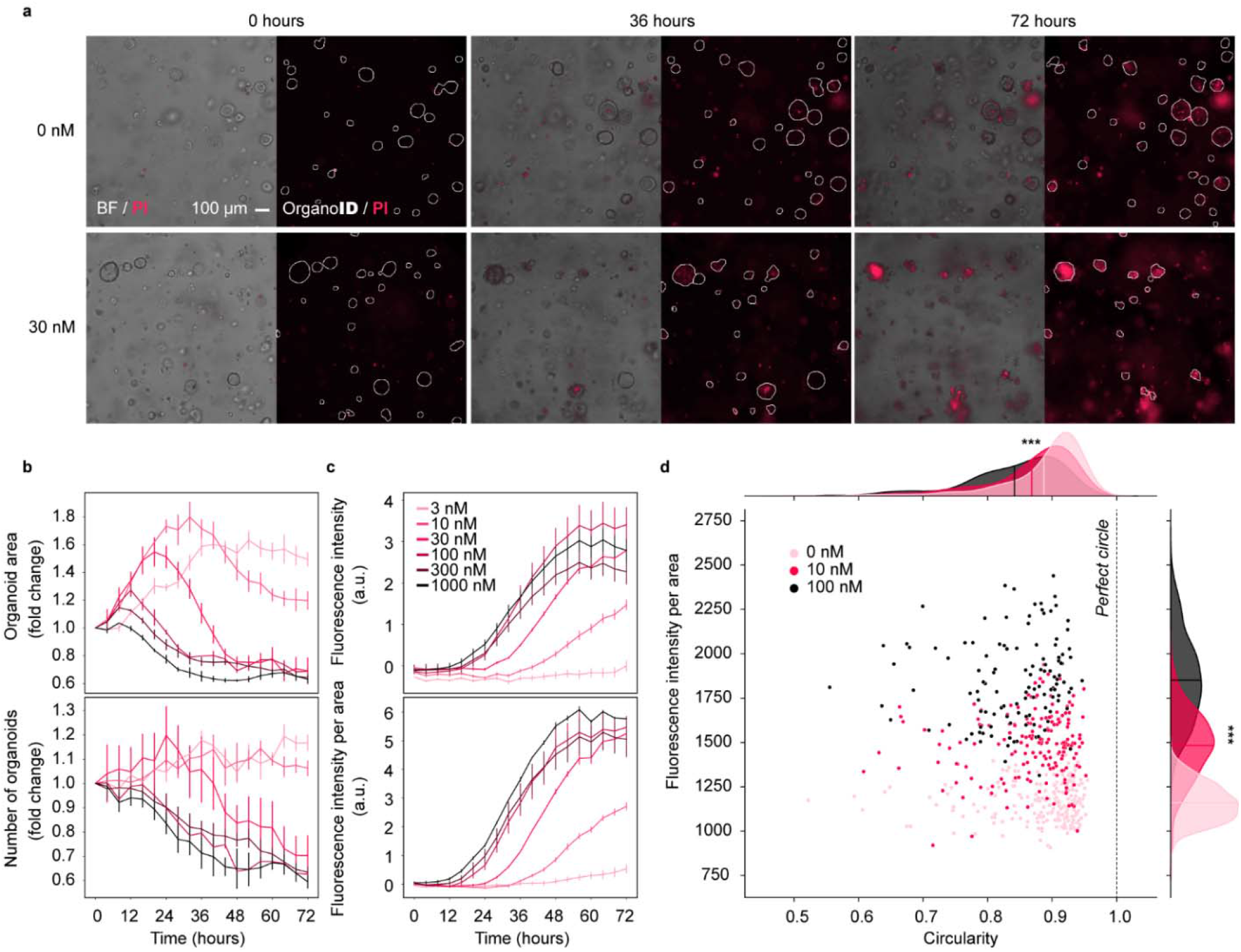
**(a)** PDAC organoids were treated with a serial dilution of gemcitabine (3 nM to 1,000 nM) and imaged over 72 hours. Propidium iodide (PI) was used to fluorescently label dead organoids. Shown are representative brightfield images from three time points for control and 30 nM gemcitabine conditions. OrganoID was used to identify organoid contours, which are displayed on top of the PI fluorescence channel. **(b)** OrganoID measurements of total organoid area (top) and number of organoids (bottom) over time for each concentration of gemcitabine. Measurements were normalized to the initial timepoint. Error bars represent standard error of the mean (n=6). **(c)** Total PI fluorescence intensity above control for each concentration of gemcitabine (top). Fluorescence intensity of OrganoID-identified regions were then summed and divided by the total organoid area (bottom), which improved separation of responses to different concentrations of gemcitabine. **(d)** Single-organoid measurements of fluorescence intensity, normalized to single-organoid area, and organoid circularity for three concentrations of gemcitabine at the 72-hour endpoint. Organoid circularity is defined as the ratio of the perimeter of a circle with the same area as the organoid to the actual perimeter of the organoid. Dots in the joint scatterplot are individual organoids. Marginal plots are kernel density estimates of the distribution of organoids for each concentration of gemcitabine. The central line for each distribution is the mean. ***: p < 0.001 for one-way ANOVA comparison of means across the three gemcitabine concentrations.

### OrganoID uncovers organoid morphology changes predictive of drug response

Changes in organoid morphology can indicate important phenotypic responses and state transitions. For example, some tumor organoid models grow into structures with invasive projections into the culture matrix, reflecting epithelial-mesenchymal transition; the addition of certain chemotherapeutic agents prevents development of these protrusions with minimal effects on overall organoid size^24^. These important responses can be investigated through single-organoid image analysis that captures the precise contour of each organoid. We used OrganoID to automatically profile the morphology of individual organoids at the endpoint of a gemcitabine dose-response experiment. Phase-contrast images of organoids exposed to control, 10 nM, and 100 nM of gemcitabine were analyzed with OrganoID to detect and identify individual organoid contours. Each organoid shape was then used to calculate the organoid circularity, a morphological measurement that compares the organoid perimeter to the perimeter of a perfect circle. This analysis uncovered that organoid circularity was significantly different across gemcitabine dosages (p<0.001), which supports disruption of organoid architecture by gemcitabine **(Figure 3d)**. This worked example demonstrates the advantages of OrganoID for automated bulk and single-organoid morphological analysis of time-course experiments without the need for live-cell fluorescence techniques.

## DISCUSSION

Organoids have revolutionized biomedical research through improved model representation of native tissues and organ systems. However, the field has yet to fully enter the high-throughput experimental space. A central bottleneck is the challenge of automated response measurement and analysis in large numbers of microscopy images. Organoids exhibit striking diversity in morphology and size and can move through their 3D environment into and out of the focal plane; current image processing tools have not quite been able to capture these aspects in a robust manner. We developed OrganoID to bridge this gap and automate the process of accurate pixel-by-pixel organoid identification and tracking over space and time.

Experimental replicates in organoid studies can differ in the number and distribution of sizes of organoids. This difference between identical conditions requires per-sample normalization of response measurements to account for differences in baseline growth of organoid colonies. There are several commercially available live-cell assays that can facilitate normalization in 2D culture. However, these same assays have proven to be difficult for organoid use due to the production of toxic photobleaching byproducts, limited diffusion through the gel matrix, and nonspecific staining of the gel matrix that results in a considerable background signal. Another available option is to genetically modify each organoid sample to express fluorescent proteins, however this increases experimental time and complexity and may alter cellular dynamics from the original tissue sample. The OrganoID platform can be leveraged for accurate normalization of standard organoid assays without live-cell fluorescence methods. OrganoID is also uniquely useful for efficient quantification of single-organoid morphological features, such as circularity, that can reflect important dynamic responses.

Additionally, we have contributed a manually segmented organoid image dataset for use in other computational platforms. OrganoID has demonstrated compatibility with organoids of various sizes, shapes, and sample concentrations as well as various optical configurations. Most excitingly, the OrganoID model was trained and validated on images of PDAC organoids but still demonstrated excellent generalization to images of other types of organoids, including those derived from colon tissue, lung tissue, and adenoid cystic carcinoma.

OrganoID was trained with and tested on a diverse, but relatively small set of images. Despite the suggested generalizability of OrganoID to various samples and optical configurations, performance may still differ with other types of organoids with significantly different morphology. As well, OrganoID can only detect and assign a single organoid to each pixel in an image. While the platform can appropriately identify contours of organoids in physical contact, it cannot distinguish organoids that overlap across the focal plane. These limitations can be overcome with additional validation, an expanded training dataset, as well as the use of multiple focal planes for image analysis.

Our image analysis platform serves as an important tool for the use of organoids as physiologically relevant models in high-throughput research. The platform can accurately capture detailed morphological measurements of individual organoids in live-cell microscopy experiments without the use of genetic modifications or potentially cytotoxic dyes. These metrics can reveal important organoid responses that might be otherwise obscured with the use of bounding-box tools. The ability of the platform to generalize to a range of organoid types without any parameter tuning also reflects the potential for the platform to standardize morphological assay readouts and improve measurement reproducibility. OrganoID achieves comprehensive and expedient image analysis of organoid experiments to enable the broader use of organoids as tissue models for high-throughput investigations into physiological and pathological phenomena.

## Supporting information

Video 1: Network training

Video 2: Single-organoid tracking

## Acknowledgements

N/A.

## Author contributions

J.M.M., B.S., S.S. Kashaf. and S.T. were involved with the conception of the platform. J.M.M. and B.S. wrote the manuscript, collaborated on platform design, and created the manual organoid segmentation dataset. B.S. performed all organoid experiments and the associated data/statistical analysis and interpretation. S.S. Kashaf developed and evaluated the initial deep learning segmentation approach. J.M.M. optimized the neural network, developed the single-organoid pipeline, and implemented the OrganoID software architecture, image processing algorithms, and performance quantification tools. R. B. and D. I. developed and provided lung organoids. C.W., E.I. and L.S. developed Adenoid Cystic Carcinoma organoids. M. Bielski and S.S. Kupfer developed and provided human colonic organoids. P. L. provided technical support with independent software and model validation. A.R., M. Bilgic, S.T. supervised the work and provided technical support. All authors commented on the manuscript.

## METHODS

### Organoid culture and image acquisition

Tumor organoid cultures derived from pancreatic adenocarcinoma (PDAC) patients were isolated and prepared as previously described^30^. PDAC organoids were cultured and imaged in a microfluidic platform^27^ or a 24-well suspension culture plate (Thermo Fisher, 144530). Human colonic organoids were cultured similarly to PDAC organoids, with Matrigel and growth or differentiation media as described in established methods^30^. Distal respiratory organoids were obtained from the Ishay-Ronen Lab at the Sheba Medical Center and cultured through a previously described protocol^31^ in 24-well plates. Images of adenoid cystic carcinoma (ACC) organoids were directly obtained from Dr. Weber and colleagues from University of Chicago Medicine. The pancreatic and airway lung organoids were cultured and imaged on an automated translational stage of an inverted microscope (Nikon Eclipse Ti) enclosed in an environmentally controlled chamber (Life Imaging Service GmbH, Basel, Switzerland). The enclosure provides temperature, humidity, and CO2 gas control to maintain adequate cell culture conditions for the organoids. Organoids were cultured at a constant 37°C, 5% CO2, a humidity flow rate of 25-30 L/hour, and 95-100% relative humidity. Images of the organoids were acquired through the standard microscope software that is capable of automatically acquiring images at different positions, *Z*-planes/stacks, and multiple fluorescent filters (NIS-Elements software, Japan). The microscope was equipped with a digital complementary metal-oxide semiconductor (CMOS) camera (ORCA-Flash 4.0, Hamamatsu, Japan), which imaged the organoids using a x10 objective at 2 to 4-hour intervals.

### Neural network design and implementation

The neural network architecture was based on the *u-net* encoder-decoder network, which has produced successful results for a variety of biomedical image segmentation tasks^**25**^. *u-net* uses several convolutional filters for each layer of neurons and doubles the number of filters per-channel for each sequential layer. The original *u-net* implementation uses 64 filters in the first layer, which results in quite a large number of trainable parameters across the full network (over 30 million). We sequentially reduced the number of filters in the first layer by powers of two to reach a minimal value that preserved performance on the validation dataset. The final OrganoID *u-net* structure uses only 8 filters in the first layer, which results in a network structure with less than 500,000 trainable parameters (a 98% reduction compared to the original implementation). This simplification reduces the computational power needed to train and use the network and, in theory, decreases network overfitting. All convolutional neurons were set to compute outputs with the exponential linear unit activation function^**26**^. The final 1×1 convolution was set with a sigmoid activation function to produce a normalized output that corresponded to the probability of an organoid at each pixel. All images are auto-contrasted and resized to 512×512 pixels before training and inference. Python was used for the entire OrganoID platform and Keras (an interface to the TensorFlow library) was used for network expression, training, and operation. The TensorFlow Lite API was used to minimize the memory footprint and number of software dependencies required for network inference in the OrganoID standalone distribution.

### Training procedure and ground-truth dataset creation

Ground-truth segmentations were created by hand with Pinta and Aseprite, two open-source image editing programs. 66 ground-truth segmentations were created from brightfield and phase-contrast microscopy images of PDAC organoids and were randomly spilt into datasets for training (52 image pairs) and validation (14 image pairs). The Augmentor Python package^32^ was used to apply random rotation, zoom, shear, elastic distortion, and skew operations to the training dataset for data augmentation. The neural network was trained with the Adam stochastic optimization algorithm^33^ at a learning rate of 0.001. Unweighted binary cross-entropy between predicted and ground-truth segmentations was used for the loss function. Layer weights were initialized with the He method^34^. An early stopping rule was used to halt training once performance on the validation dataset reached a minimum (i.e. once 10 epochs pass with no improvement in validation loss). The batch size was set to train on 8 augmented images for every round of backpropagation. Dropout regularization was introduced after all convolutions to randomly set 12.5% of neuron weights to zero after each batch. After each epoch, a copy of the model was saved for additional evaluation of the training process.

### Dataset for evaluation of network performance and generalizability

The network structure and training process were manually tuned for performance on the PDAC validation dataset. After optimization, we locked out any further changes to all hyperparameters and evaluated final network performance and generalizability on a separate dataset of 28 organoid microscopy images. The dataset included images of organoids derived from PDAC (10), benign colon tumor cells (6), lung epithelia (6), and salivary adenoid cystic carcinoma (6). The images were obtained through brightfield and phase-contrast microscopy from multiple microscopy cores and manually labeled by two independent reviewers.

### Single-organoid identification and segmentation refinement

The *scipy*^35^ and *scikit-image*^36^ packages were used to identify individual organoids from the network detection images. Detection images were thresholded at 0.5 and passed through a morphological opening operation to remove weak and noisy predictions. Partial derivatives were computed with a Sobel filter, passed through a Gaussian filter (s=2), and converted to an edge mask with a hysteresis threshold filter (T_hi_=0.05, T_lo_=0.005), which was then was used to mark organoid centers. The watershed method was used to identify filled organoid contours, with the organoid centers as label initializers and the network prediction image as an inverted heightmap. Labeled organoids were morphologically filled and discarded if the total area was below 200 pixels. Images were morphologically processed with *scikit-image* to record properties of identified organoids.

### Tracking of individual organoids over time

The tracking algorithm builds a cost-assignment matrix for each image in a time-lapse sequence, where each row corresponds to an organoid from the previous image and each column corresponds to an organoid in the next image. Each matrix entry is the cost of assigning a given organoid detection in the current image to a detection in the previous image. The cost function was defined as the inverse of the number of shared pixels between the two organoids. As such, the assignment cost between two detections will be minimal for those that are of similar shape and location. The cost-assignment matrix is also padded with additional rows and columns to allow for “ pseudo-assignments” that represent missing or newly detected organoids. Finally, the Munkres variant of the Hungarian algorithm^29^ was used to minimize the cost-assignment matrix to find an optimal matching between organoid detections in the previous image and the detections in the current image

### Statistical methods for platform validation

Statistical analysis was performed with the *numpy* and *scipy* packages in Python. Network performance on the testing dataset was evaluated as the pixel-wise intersection-over-union (IOU) of predictions compared to ground-truth segmentations. A single IOU value was computed for each prediction/ground-truth pair and summarized as a mean IOU and standard deviation over the dataset. For agreement of single-organoid counts and measurements, the Lin concordance correlation coefficient (CCC) was computed as

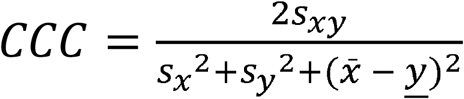

The Fisher transformation (*artanh(CCC)*) and the inverse Fisher transformation (*tanh(z*_*LO*_, *Z*_*HI*_)) were also used to obtain a z-value and construct a 95% confidence interval for CCC statistics. All plots were generated with the *matplotlib* and *seaborn* packages in Python.

### Drug screening experiments

PDAC organoids were grown for one week in standard culture conditions and then treated with gemcitabine hydrochloride (G6423, Sigma) at six serial dilutions from 3 nM to 1000 nM with negative control. Propidium iodide (Thermo Fisher P3566) was used to fluorescently measure cellular death and relative viability of organoids in real-time at 4-hour intervals over 72 hours. An MTS proliferation assay (Promega, G3580) was also performed at the end of the experiment to determine cell viability for each condition according to manufacturer’s instructions. Statistical hypothesis testing was performed with *scipy* and MATLAB. Organoid circularity was computed as the circumference of a circle with equivalent area divided by the actual perimeter of the organoid.

### Validation rigor and indications of robustness

The OrganoID neural network was tested on images of four types of organoids taken at multiple microscopy cores. Images were manually segmented by two independent evaluators. OrganoID single-organoid count and area measurements were compared against the corresponding measurements of the manual test segmentations. OrganoID tracking results were compared to manually tracked images. The OrganoID platform and test microscopy images were distributed to an independent user to confirm reproducibility of all metrics, statistics, and figures.

### Platform availability

We have released the OrganoID platform open-source and freely licensed on GitHub (https://github.com/jono-m/OrganoID). The repository includes all source code and a compiled executable, as well as the entire training and testing dataset, usage instructions, and scripts used for the examples presented in this paper. The network training module is also included on the repository to allow further model training to improve performance for any untested applications.

## References

1. Kretzschmar, K. & Clevers, H. Organoids: Modeling Development and the Stem Cell Niche in a Dish. Developmental Cell 38, 590–600 (2016).

2. Driehuis, E. et al. Oral Mucosal Organoids as a Potential Platform for Personalized Cancer Therapy. Cancer Discovery 9, 852–871 (2019).

3. Dutta, D., Heo, I. & Clevers, H. Disease Modeling in Stem Cell-Derived 3D Organoid Systems. Trends in Molecular Medicine 23, 393–410 (2017).

4. Sachs, N. et al. A Living Biobank of Breast Cancer Organoids Captures Disease Heterogeneity. Cell 172, 373-386.e10 (2018).

5. Clevers, H. & Tuveson, D. A. Organoid Models for Cancer Research. Annual Review of Cancer Biology 3, 223–234 (2019).

6. Garreta, E. et al. Rethinking organoid technology through bioengineering. Nature Materials 20, 145–155 (2020).

7. Pettinato, G. et al. Spectroscopic label-free microscopy of changes in live cell chromatin and biochemical composition in transplantable organoids. Sci Adv 7, eabj2800 (2021).

8. Mead, B. E. et al. Screening for modulators of the cellular composition of gut epithelia via organoid models of intestinal stem cell differentiation. Nature Biomedical Engineering (2022) doi:10.1038/s41551-022-00863-9.

9. Brandenberg, N. et al. High-throughput automated organoid culture via stem-cell aggregation in microcavity arrays. Nat Biomed Eng 4, 863–874 (2020).

10. Zhou, Z. et al. An organoid-based screen for epigenetic inhibitors that stimulate antigen presentation and potentiate T-cell-mediated cytotoxicity. Nat Biomed Eng 5, 1320–1335 (2021).

11. Wiedenmann, S. et al. Single-cell-resolved differentiation of human induced pluripotent stem cells into pancreatic duct-like organoids on a microwell chip. Nat Biomed Eng 5, 897–913 (2021).

12. Richards, D. J. et al. Human cardiac organoids for the modelling of myocardial infarction and drug cardiotoxicity. Nat Biomed Eng 4, 446–462 (2020).

13. Tanaka, N. et al. Three-dimensional single-cell imaging for the analysis of RNA and protein expression in intact tumour biopsies. Nat Biomed Eng 4, 875–888 (2020).

14. Kim, S. et al. Comparison of Cell and Organoid-Level Analysis of Patient-Derived 3D Organoids to Evaluate Tumor Cell Growth Dynamics and Drug Response. SLAS Discov 25, 744–754 (2020).

15. Dekkers, J. F. et al. High-resolution 3D imaging of fixed and cleared organoids. Nature Protocols 14, 1756–1771 (2019).

16. Hof, L. et al. Long-term live imaging and multiscale analysis identify heterogeneity and core principles of epithelial organoid morphogenesis. BMC Biol 19, 37–37 (2021).

17. Bailey, S. R. & Maus, M. V. Gene editing for immune cell therapies. Nature Biotechnology 37, 1425–1434 (2019).

18. Ang, L. T. et al. A Roadmap for Human Liver Differentiation from Pluripotent Stem Cells. Cell Rep 22, 2190–2205 (2018).

19. Riss, T. & Trask, O. J., Jr. Factors to consider when interrogating 3D culture models with plate readers or automated microscopes. In Vitro Cell Dev Biol Anim 57, 238–256 (2021).

20. Borten, M. A., Bajikar, S. S., Sasaki, N., Clevers, H. & Janes, K. A. Automated brightfield morphometry of 3D organoid populations by OrganoSeg. Sci Rep 8, 5319–5319 (2018).

21. Kassis, T., Hernandez-Gordillo, V., Langer, R. & Griffith, L. G. OrgaQuant: Human Intestinal Organoid Localization and Quantification Using Deep Convolutional Neural Networks. Sci Rep 9, 12479–12479 (2019).

22. Kok, R. N. U. et al. OrganoidTracker: Efficient cell tracking using machine learning and manual error correction. PLoS One 15, e0240802–e0240802 (2020).

23. Larsen, B. M. et al. A pan-cancer organoid platform for precision medicine. Cell Reports 36, 109429 (2021).

24. Zhao, N. et al. Morphological screening of mesenchymal mammary tumor organoids to identify drugs that reverse epithelial-mesenchymal transition. Nat Commun 12, 4262–4262 (2021).

25. Ronneberger, O., Fischer, P. & Brox, T. U-Net: Convolutional Networks for Biomedical Image Segmentation. Lecture Notes in Computer Science 234–241 (2015) doi:10.1007/978-3-319-24574-4_28.

26. Clevert, D.-A., Unterthiner, T. & Hochreiter, S. Fast and Accurate Deep Network Learning by Exponential Linear Units (ELUs). (2015) doi:10.48550/ARXIV.1511.07289.

27. Schuster, B. et al. Automated microfluidic platform for dynamic and combinatorial drug screening of tumor organoids. Nat Commun 11, 5271–5271 (2020).

28. Canny, J. A Computational Approach to Edge Detection. IEEE Transactions on Pattern Analysis and Machine Intelligence PAMI-8, 679–698 (1986).

29. Munkres, J. Algorithms for the Assignment and Transportation Problems. Journal of the Society for Industrial and Applied Mathematics 5, 32–38 (1957).

30. Romero-Calvo, I. et al. Human Organoids Share Structural and Genetic Features with Primary Pancreatic Adenocarcinoma Tumors. Mol Cancer Res 17, 70–83 (2019).

31. Sachs, N. et al. Long-term expanding human airway organoids for disease modeling. EMBO J 38, e100300 (2019).

32. D Bloice, M., Stocker, C. & Holzinger, A. Augmentor: An Image Augmentation Library for Machine Learning. The Journal of Open Source Software 2, 432 (2017).

33. Kingma, D. P. & Ba, J. Adam: A Method for Stochastic Optimization. (2014) doi:10.48550/ARXIV.1412.6980.

34. He, K., Zhang, X., Ren, S. & Sun, J. Delving Deep into Rectifiers: Surpassing Human-Level Performance on ImageNet Classification. in (IEEE, 2015). doi:10.1109/iccv.2015.123.

35. Virtanen, P. et al. SciPy 1.0: fundamental algorithms for scientific computing in Python. Nat Methods 17, 261–272 (2020).

36. van der Walt, S. et al. scikit-image: image processing in Python. PeerJ 2, e453–e453 (2014).

