## Supplemental Information for "OrganoID: a versatile deep learning platform for tracking and analysis of single-organoid dynamics"

**Videos:**
**V1:** Network training at the end of each epoch: representative images
**V2:** Organoid tracking video

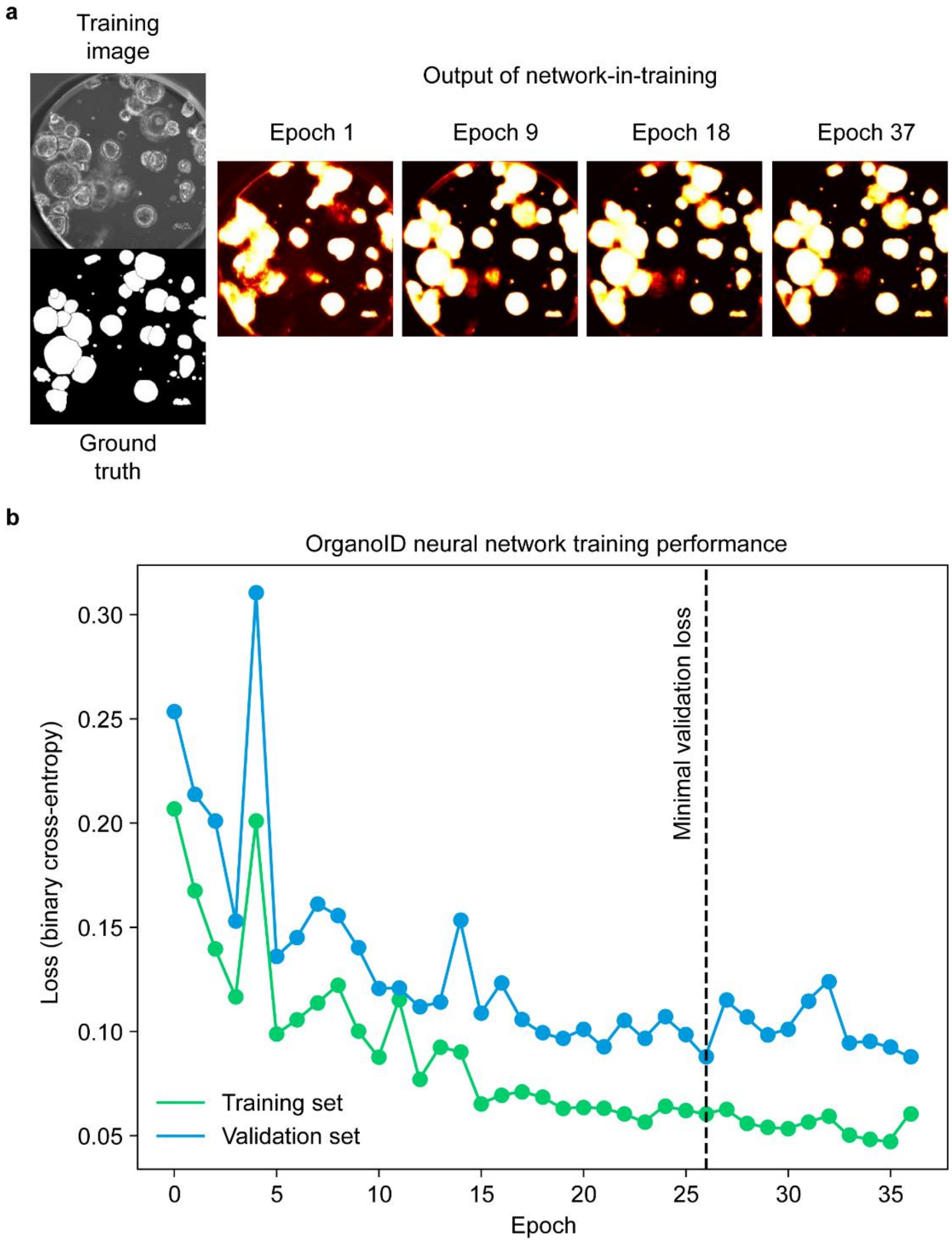


**Supplementary Figure 1. Network training performance.** (a) The OrganoID neural network predicts the probability that an organoid is present at each pixel. Shown are network predictions produced by intermediate models at selected epochs through the training process. (b) Network training was stopped after 37 epochs, once a minimum binary cross-entropy loss on the validation dataset was reached

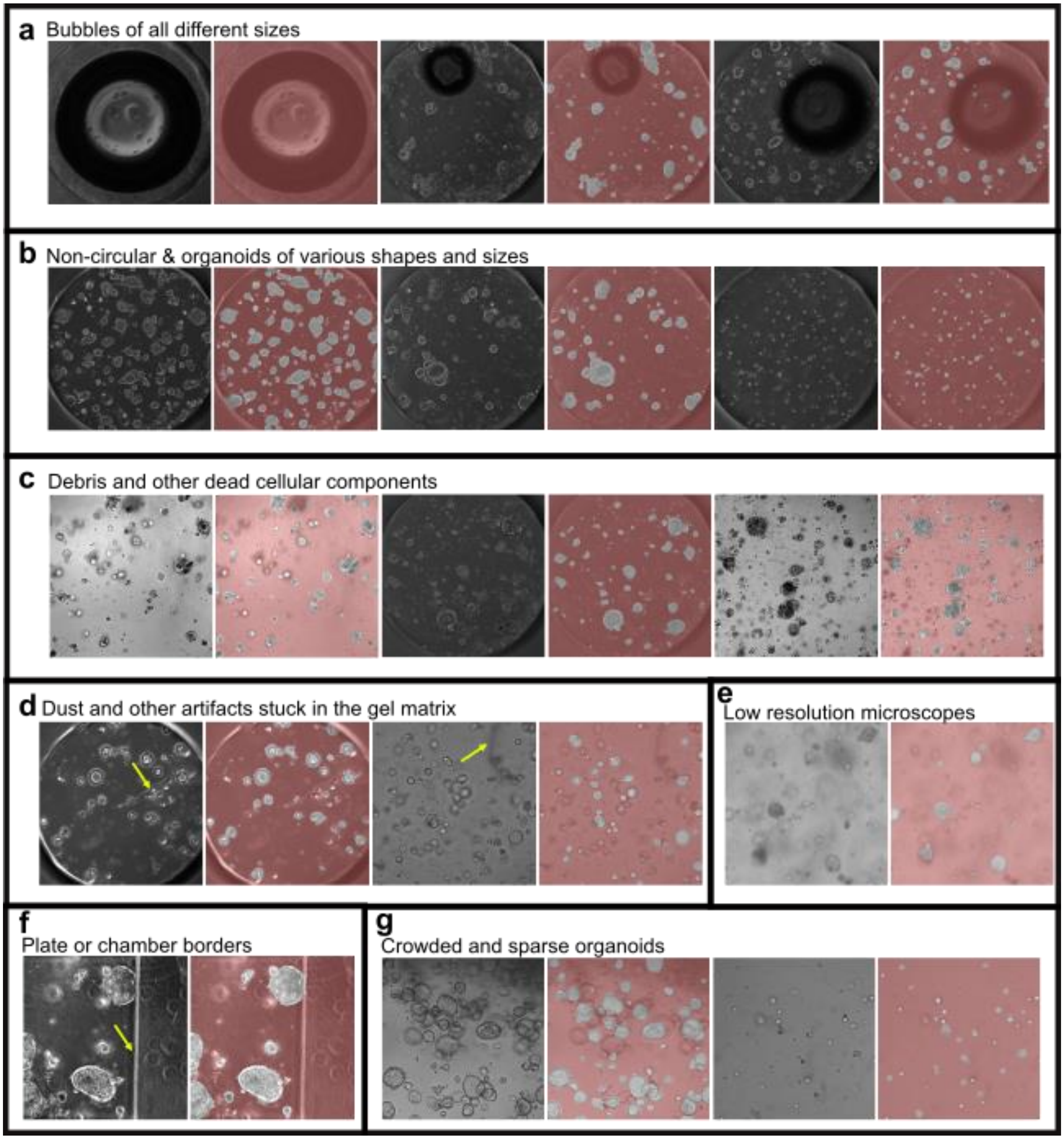


**Supplementary Figure 2. Exclusion of non-organoid artifacts.** OrganoID ignored bubbles (a), debris (c-d), and plate or microfluidic chamber borders (f) to accurately identify organoids that exhibit diverse morphology and sizes, even within a single sample (b). OrganoID can also handle various optical configurations, including low-resolution or poorly-lit images (e). Gel droplets can support densely-packed or isolated organoids, which can all be detected with OrganoID (g).

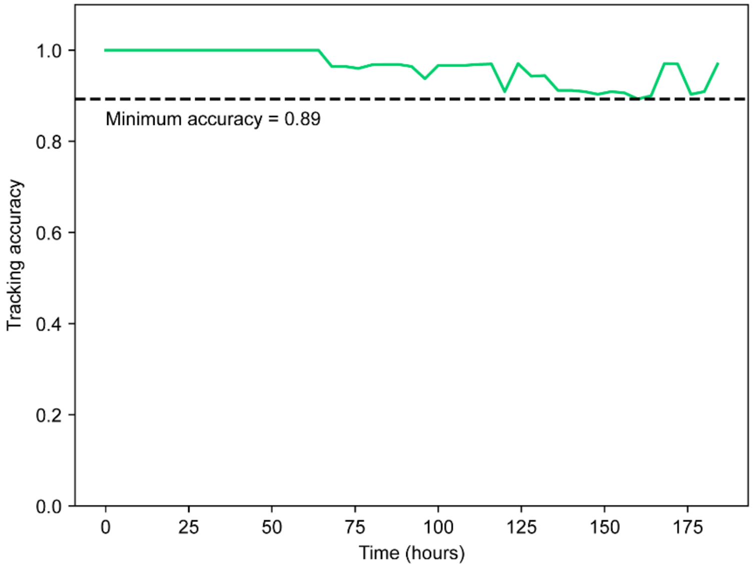


**Supplementary Figure 3. Tracking accuracy over time.** A time-lapse microscopy experiment was analyzed with OrganoID to identify organoids in each image. OrganoID was then used to match identified organoids across frames to build single-organoid tracks. The identified organoids were also matched by hand to assess tracking performance. Accuracy was defined the number of organoid track labels in agreement divided by the total number of organoids present at each frame.

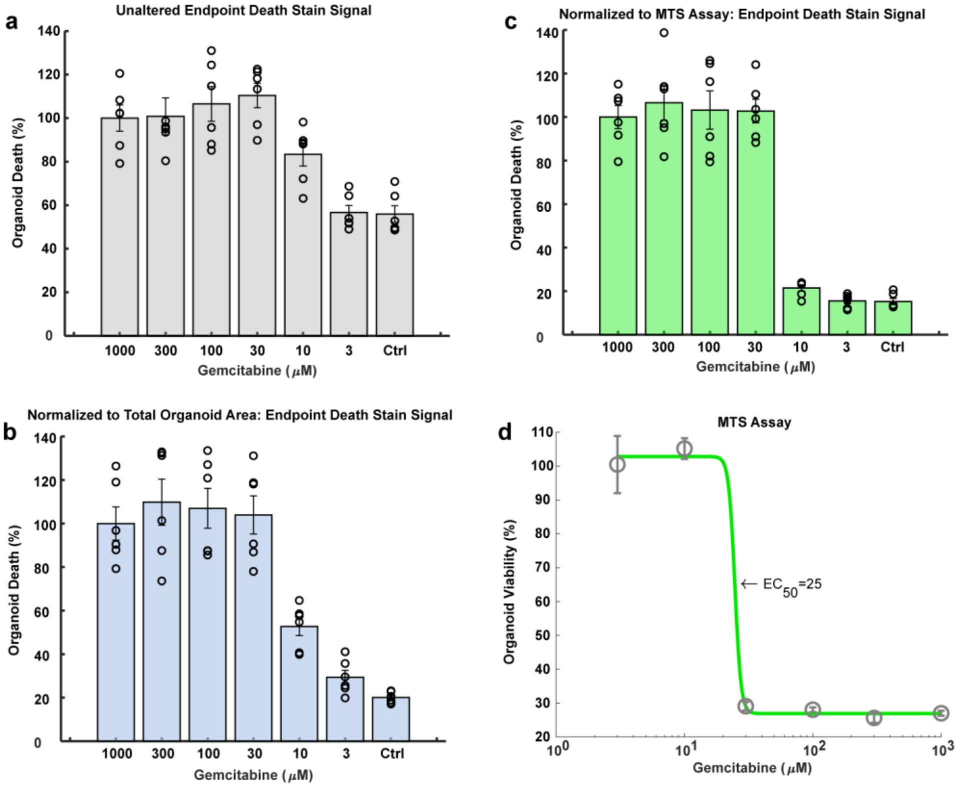


**Supplementary Figure 4. Comparison of drug treatment data analysis normalized with OrganoID.** (a) Intensity sum of PI fluorescence at the end of the 72-hour drug treatment period for each concentration of gemcitabine. (b) Intensity sum of PI fluorescence divided by the total organoid area, computed with OrganoID, in each sample at the 72-hour time point, which normalizes for differences in the baseline number and sizes of organoids in each replicate. (c) A gold standard endpoint proliferation assay, MTS, was conducted to measure total organoid viability in each sample. Here, the intensity sum of PI fluorescence was instead normalized with the MTS readout for comparison to OrganoID normalization. All PI measurements were normalized to the readout from the strongest drug concentration (1000 nM) and presented as mean values ± SEM, n = 6. One-Way ANOVA statistical results in Table S1. (d) A viability curve was computed from the MTS assay and demonstrated a half-maximal effective concentration (EC50) of 25 μM.

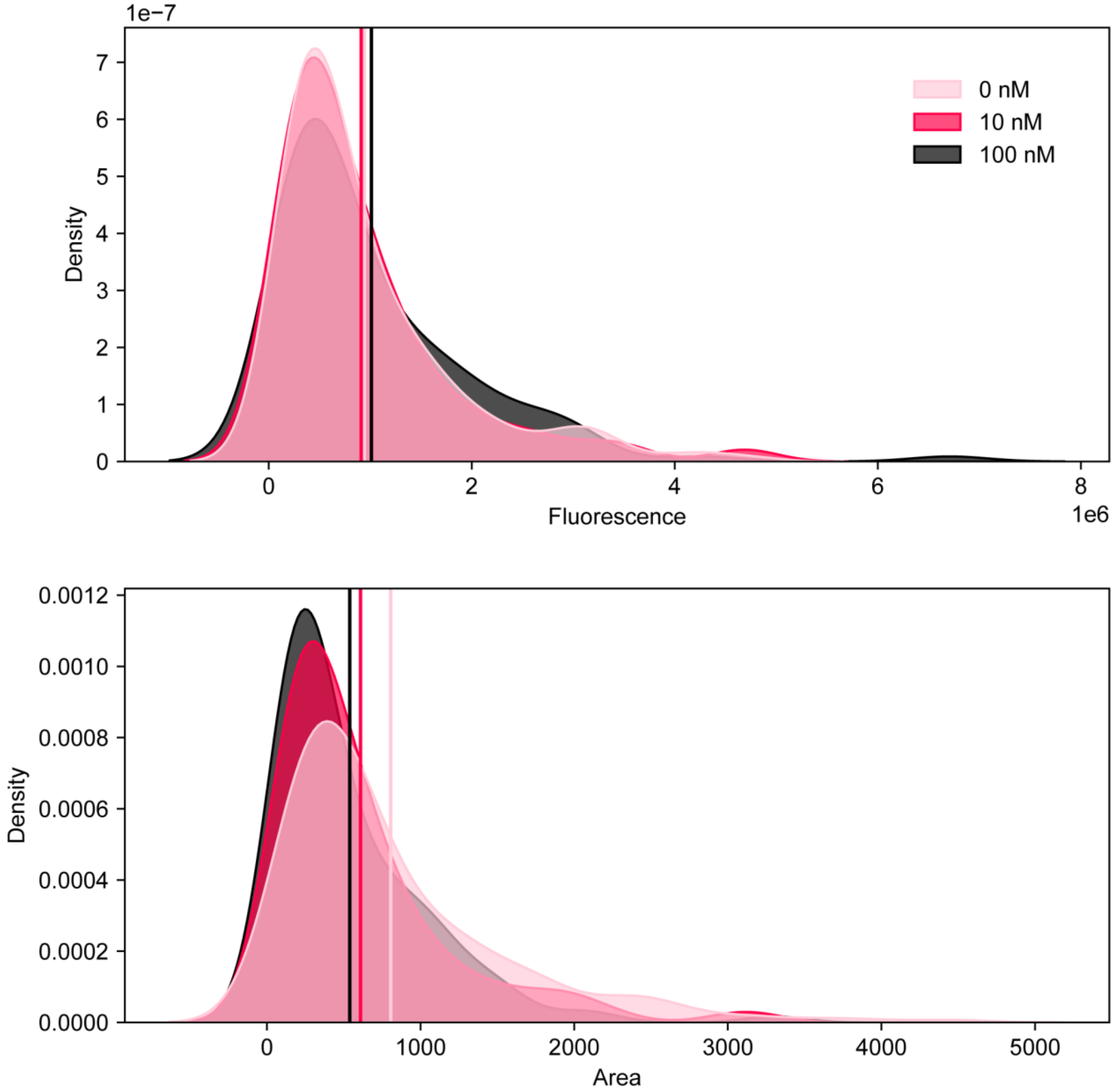


**Supplementary Figure 5. Single-organoid fluorescence and area distribution.** Kernel density estimates for the distribution of single-organoid fluorescence and area at the endpoint of a 72-hour gemcitabine dose-response experiment. Area is measured in pixels at 6.8644 $\mu m^2$/pixel.

**Supplementary Table 1: One-Way ANOVA Calculated P-Values (P<0.001)**

| Condition 1 | Condition 2 | PI Data | PI Data Normalized with Area | PI Data Normalized with MTS |
|---|---|---|---|---|
| **1000nM** | **300nM** | 1 | 0.956628 | 0.975909 |
| **1000nM** | **100nM** | 0.987028 | 0.992248 | 0.999517 |
| **1000nM** | **30nM** | 0.884143 | 0.999688 | 0.999789 |
| **1000nM** | **1 nM** | 0.465339 | 0.0008 | 3.72E-08 |
| **1000nM** | **3nM** | 0.000246 | 8.84E-07 | 3.71E-08 |
| **1000nM** | **Ctrl** | 0.000191 | 9.51E-08 | 3.71E-08 |
| **300nM** | **100nM** | 0.993637 | 0.999953 | 0.999353 |
| **300nM** | **30nM** | 0.918481 | 0.996913 | 0.998727 |
| **300nM** | **1 nM** | 0.407788 | 4.48E-05 | 3.71E-08 |
| **300nM** | **3nM** | 0.000185 | 8.68E-08 | 3.71E-08 |
| **300nM** | **Ctrl** | 0.000143 | 4.08E-08 | 3.71E-08 |
| **100nM** | **30nM** | 0.999275 | 0.999928 | 1 |
| **100nM** | **1 nM** | 0.125988 | 0.000104 | 3.71E-08 |
| **100nM** | **3nM** | 2.52E-05 | 1.49E-07 | 3.71E-08 |
| **100nM** | **Ctrl** | 1.95E-05 | 4.52E-08 | 3.71E-08 |
| **30nM** | **1 nM** | 0.04618 | 0.000255 | 3.71E-08 |
| **30nM** | **3nM** | 6.55E-06 | 3.07E-07 | 3.71E-08 |
| **30nM** | **Ctrl** | 5.08E-06 | 5.62E-08 | 3.71E-08 |
| **10nM** | **3nM** | 0.050677 | 0.272317 | 0.985243 |
| **10nM** | **Ctrl** | 0.041342 | 0.040022 | 0.982259 |
| **3nM** | **Ctrl** | 1 | 0.966961 | 1 |

**Appendix 1: Training, Validation and Testing Image Index**

| Training & Validation | | | | |
|---|---|---|---|---|
| **Label** | **Experiment - Date** | **Brightfield (BF) or Phase Contrast (PC)** | **Microfluidic or Plate Experiment** | **PDAC Patient #** |
| 0 | 1-22-2018 | PC | Microfluidic | 2 |
| 1 | 1-22-2018 | PC | Microfluidic | 2 |
| 2 | 1-22-2018 | PC | Microfluidic | 2 |
| 3 | 1-22-2018 | PC | Microfluidic | 2 |
| 4 | 1-22-2018 | PC | Microfluidic | 2 |
| 5 | 1-22-2018 | PC | Microfluidic | 2 |
| 6 | 1-22-2018 | PC | Microfluidic | 2 |
| 7 | 1-22-2018 | PC | Microfluidic | 2 |
| 8 | 1-22-2018 | PC | Microfluidic | 2 |
| 9 | 1-22-2018 | PC | Microfluidic | 2 |
| 10 | 1-22-2018 | PC | Microfluidic | 2 |
| 11 | 1-22-2018 | PC | Microfluidic | 2 |
| 12 | 1-22-2018 | PC | Microfluidic | 2 |
| 13 | 1-22-2018 | PC | Microfluidic | 2 |
| 14 | 1-22-2018 | PC | Microfluidic | 2 |
| 15 | 1-22-2018 | PC | Microfluidic | 2 |
| 16 | 1-22-2018 | PC | Microfluidic | 2 |
| 17 | 1-22-2018 | PC | Microfluidic | 2 |
| 18 | 1-22-2018 | PC | Microfluidic | 2 |
| 19 | 1-22-2018 | PC | Microfluidic | 2 |
| 20 | 1-22-2018 | PC | Microfluidic | 2 |
| 21 | 1-22-2018 | PC | Microfluidic | 2 |
| 22 | 1-22-2018 | PC | Microfluidic | 2 |
| 23 | 1-22-2018 | PC | Microfluidic | 2 |

| | | | | |
|---|---|---|---|---|
| 24 | 1-22-2018 | PC | Microfluidic | 2 |
| 25 | 1-22-2018 | PC | Microfluidic | 2 |
| 26 | 1-22-2018 | PC | Microfluidic | 2 |
| 27 | 1-22-2018 | PC | Microfluidic | 2 |
| 28 | 1-22-2018 | PC | Microfluidic | 2 |
| 29 | 1-22-2018 | PC | Microfluidic | 2 |
| 30 | 1-22-2018 | PC | Microfluidic | 2 |
| 31 | 1-22-2018 | PC | Microfluidic | 2 |
| 32 | 2-14-2021 | BF | Plate | 1 |
| 33 | 2-14-2021 | BF | Plate | 1 |
| 34 | 7-16-2018 | PC | Microfluidic | 1 |
| 35 | 7-16-2018 | PC | Microfluidic | 1 |
| 36 | 7-16-2018 | PC | Microfluidic | 1 |
| 37 | 5-12-2017 | PC | Microfluidic, V1 | 2 |
| 38 | 7-16-2018 | PC | Microfluidic | 1 |
| 39 | 1-22-2018 | PC | Microfluidic | 2 |
| 40 | 1-22-2018 | PC | Microfluidic | 2 |
| 41 | 1-22-2018 | PC | Microfluidic | 2 |
| 42 | 6-22-2018 | PC | Plate | 2 |
| 43 | 6-22-2018 | PC | Plate | 2 |
| 44 | 6-22-2018 | PC | Plate | 2 |
| 45 | 3-15-2018 | PC | Microfluidic | 1 |
| 46 | 3-25-2021 | BF | Plate | 2 |
| 47 | 3-25-2021 | BF | Plate | 2 |
| 48 | 3-25-2021 | BF | Plate | 2 |
| 49 | 3-15-2018 | PC | Microfluidic | 1 |
| 50 | 3-25-2021 | BF | Plate | 1 |
| 51 | 3-25-2021 | BF | Plate | 1 |

| | | | | |
|---|---|---|---|---|
| 52 | 3-25-2021 | BF | Plate | 1 |
| 53 | 3-06-2019 | PC | Microfluidic | 2 |
| 54 | 3-06-2019 | PC | Microfluidic | 2 |
| 55 | 3-25-2021 | PC | Plate | 2 |
| 56 | 3-28-2019 | PC | Microfluidic | 2 |
| 57 | 3-28-2019 | PC | Microfluidic | 2 |
| 58 | 3-15-2018 | PC | Microfluidic | 1 |
| 59 | 3-15-2018 | PC | Microfluidic | 1 |
| 60 | 3-25-2021 | BF | Plate | 2 |
| 61 | 3-25-2021 | BF | Plate | 1 |
| 62 | 3-15-2018 | PC | Microfluidics | 1 |
| 63 | 3-15-2018 | PC | Microfluidics | 1 |
| 64 | 3-28-2019 | PC | Microfluidics | 1 |
| 65 | 3-28-2019 | PC | Microfluidics | 1 |

| **Testing Images** | | | | |
|---|---|---|---|---|
| **Label** | **Experiment - Date** | **Brightfield (BF) or Phase Contrast (PC)** | **Microfluidic or Plate Experiment** | **PDAC Patient #** |
| T1 | 11-05-2018 | PC | Microfluidic | 1 |
| T2 | 1-27-2021 | BF | Plate | 1 |
| T3 | 1-27-2021 | PC | plate | 1 |
| T4 | 1-27-2021 | BF | Plate | 1 |
| T5 | 11-05-2018 | BF | Plate | 2 |
| T6 | 11-05-2018 | BF | Plate | 1 |
| T7 | 1-27-2021 | PC | Plate | 1 |
| T8 | 6-08-2018 | PC | Plate | 1 |
| T9 | 10-2020 | PC | Microfluidic | 1 |
| T10 | 3-28-2019 | PC | Microfluidic | 3 |

| Airway Organoid Testing Images | | | | |
|---|---|---|---|---|
| **Label** | **Experiment – Date** | **Brightfield (BF) or Phase Contrast (PC)** | **Microfluidic or Plate Experiment** | **Airway Patient** |
| A1 | 2-08-2021 | BF | plate | AW2 |
| A2 | 2-11-2021 | BF | plate | AW1 |
| A3 | 2-11-2021 | PC | plate | AW1 |
| A4 | 2-08-2021 | BF | plate | AW2 |
| A5 | 2-08-2021 | BF | plate | AW2 |
| A6 | 2-11-2021 | BF | plate | AW2 |

| Colon Organoid Testing Images | | | | |
|---|---|---|---|---|
| **Label** | **Experiment - Date** | **Brightfield (BF) or Phase Contrast (PC)** | **Microfluidic or Plate Experiment** | **Patient** |
| C1 | 4-21-2021 | BF | plate | N/A |
| C2 | 4-21-2021 | BF | plate | N/A |
| C3 | 4-21-2021 | BF | plate | N/A |
| C4 | 4-21-2021 | BF | plate | N/A |
| C5 | 4-21-2021 | BF | plate | N/A |
| C6 | 4-21-2021 | BF | plate | N/A |

| ACC Organoid Testing Images | | | | |
|---|---|---|---|---|
| **New Label** | **Experiment - Date** | **Brightfield (BF) or Phase Contrast (PC)** | **Microfluidic or Plate Experiment** | **Patient** |
| ACC1 | 10-22-2020 | BF | plate | N/A |
| ACC2 | 10-22-2020 | BF | plate | N/A |
| ACC3 | 10-23-2020 | BF | plate | N/A |
| ACC4 | 10-23-2020 | BF | plate | N/A |
| ACC5 | N/A | BF | plate | N/A |
| ACC6 | N/A | BF | plate | N/A |
