## Supplementary figures and images for "OrganoID: a versatile deep learning platform for tracking and analysis of single-organoid dynamics"

### Video 2: Single-organoid tracking

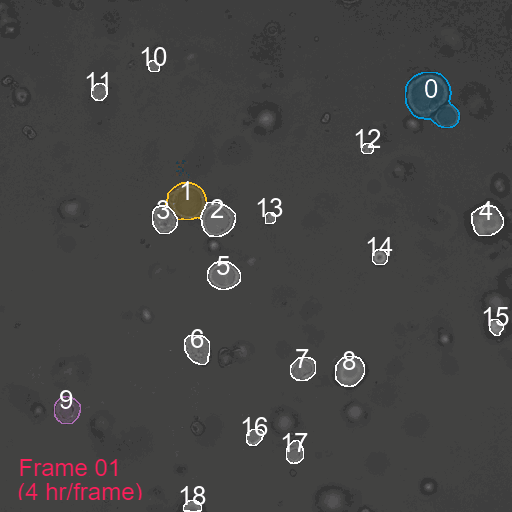
